# Investigating the Role of *Onchocerca ochengi* in Epilepsy Development: A Gerbil Model Study

**DOI:** 10.1101/2025.02.10.637387

**Authors:** Rene Bilingwe Ayiseh, Fobang Ulrick Anangafack, Judith Christine Etaka, Gamua Stanley Dobgima, Chrysantus Njobinkir Bimela, Stephen Mbigha Ghogomu, Fidelis Cho-Ngwa

**Affiliations:** Drugs and Molecular Diagnostic Laboratory (DMD), Biotechnology Unit, University of Buea, South West Region, Cameroon; Department of Biochemistry and Molecular Biology, Faculty of Science, University of Buea, South West Region, Cameroon; Molecular and Cell Biology Laboratory (MCBL), Biotechnology Unit, University of Buea, South West Region, Cameroon

**Keywords:** onchocerciasis-associated epilepsy, onchocerciasis, gerbil model, *Onchocerca ochengi*

## Abstract

**Background:** *Onchocerca volvulus* infection is linked to onchocerciasis-associated epilepsy (OAE) in humans, but the role of *Onchocerca ochengi* in epilepsy development remains unexplored. This study aimed to investigate whether *O. ochengi* infection contributes to epilepsy development.

**Methodology/Principal Findings:** Gerbils were implanted with *O. ochengi* worm masses (test group) or underwent sham surgery (control group). Behavioral and physical assessments were performed between days 15–19 using multiple tests, including the elevated plus maze, open-field, object recognition, and hanging wire tests. On day 21, gerbils were sacrificed, and body/organ weights were recorded, along with worm mass survival. Implantation of 15 worm masses resulted in 100% mortality in the test group, while implantation of 10 worm masses resulted in 53.3% mortality, with all control animals surviving. At day 21, worm mass survival averaged 1.4 out of 10, with a viability score of 93.3%. Test animals showed significant reductions in body weight and increased spleen weight compared to controls, but no significant behavioral differences were observed.

**Conclusions/Significance:** While *O. ochengi* infection caused notable physical effects, including high mortality and changes in body/organ weights, no behavioral evidence of epilepsy was observed. The high mortality rate and limited observation period restrict the interpretation of these findings. Further studies with larger cohorts and longer observation periods are needed to assess the potential role of *Onchocerca spp*. in epilepsy development. To the best of our knowledge, this study represents the first attempt to establish an animal model for OAE.

**Author Summary:** *Onchocerca volvulus* infection is linked to onchocerciasis-associated epilepsy (OAE) in humans, but we know less about whether other types of *Onchocerca*, such as *Onchocerca ochengi*, might also contribute to epilepsy. Our study aimed to investigate this question by infecting gerbils with *O. ochengi* worm masses and observing the impact on their behavior and physical health. We found that while the infection caused significant physical changes, including high mortality rates and changes in body and organ weights, there were no signs of behavioral changes typical of epilepsy. In particular, we did not see any of the usual neurological symptoms that might indicate epilepsy. These results suggest that while *O. ochengi* can affect animal health in some ways, it might not be directly involved in causing epilepsy. However, the high mortality rate in the infected gerbils and the relatively short duration of the study mean that we cannot draw firm conclusions. Future research with more animals and a longer time frame will be important to better understand whether *Onchocerca* worms contribute to epilepsy development in humans.

## Introduction

Epilepsy is a prevalent neurological disorder characterized by recurrent seizures, affecting approximately 50 million people worldwide.[1] While the causes of epilepsy are multifaceted, parasitic infections have been identified as significant contributors to seizure disorders, particularly in endemic regions. One such infection, onchocerciasis, caused by the parasitic nematode *Onchocerca volvulus*, has been linked to epilepsy in various epidemiological studies, particularly in sub-Saharan Africa.[2] This condition, referred to as onchocerciasis-associated epilepsy (OAE), has raised concerns about the potential role of the parasitic infection in triggering epilepsy. However, despite growing evidence, no direct causal relationship between onchocerciasis and epilepsy has been experimentally confirmed.

Onchocerciasis, commonly known as “river blindness,” is primarily associated with dermatological and ocular complications due to the presence of microfilariae in the skin and eyes.[3] Recent epidemiological studies, however, have suggested an alarming association between onchocerciasis and increased epilepsy incidence in high-burden areas, particularly in regions where mass treatment programs with ivermectin, the primary drug for onchocerciasis control, are insufficient.[4] While these studies point to a potential link between onchocerciasis and epilepsy, the underlying mechanisms remain unclear. Recent research has suggested that the role of *Onchocerca* infections in epilepsy could be direct or mediated indirectly through the microbiome, including viruses such as OVRV1, which may contribute to epileptogenesis.[5, 6] Further investigation is needed to clarify these potential pathways.

Given the ethical and logistical limitations of human studies, animal models serve as a crucial tool for exploring disease mechanisms. Gerbils (*Meriones unguiculatus*) are particularly useful in studying epilepsy due to their predisposition to seizure activity, making them an appropriate model for evaluating the potential epileptogenic effects of parasitic infections.[7]

The present study aims to bridge the gap in knowledge by using a gerbil model to investigate whether infection with *Onchocerca ochengi*, a close relative of *O. volvulus[8]* and a known cattle parasite, can directly induce epilepsy-like symptoms. in gerbils. Although *O. ochengi* is not directly responsible for human onchocerciasis, its close relationship to *O. volvulus* warrants exploration into whether it can contribute to neurological disturbances. This investigation addresses the absence of experimental studies linking onchocerciasis to epilepsy and explores the biological effects of *O. ochengi* worm masses on gerbil survival, behavior, and physiology.

In this study, gerbils were surgically implanted with *O. ochengi* worm masses to mimic infection, with a control group undergoing sham operations. Behavioral tests, including the elevated plus maze and open-field tests, were conducted to evaluate anxiety-like behavior and general locomotor activity, while object recognition and hanging wire tests assessed cognitive function and muscle strength, respectively. The choice of these behavioral tests aimed to identify any subtle neurological changes indicative of epilepsy or cognitive impairment, as seen in human epilepsy cases.[9]

The significance of this research lies in its potential to clarify the relationship between parasitic infections and epilepsy. Understanding the mechanisms underlying onchocerciasis-associated epilepsy could lead to better diagnostic tools and intervention strategies for reducing epilepsy prevalence in onchocerciasis-endemic regions. This study represents a valuable step toward understanding the role of O. ochengi in epilepsy development and the broader implications of parasitic infections in neuroinflammatory diseases.

## Methods

### Experimental Design and Procedure

This study involved a laboratory-based investigation using Mongolian gerbils (*Meriones unguiculatus*) as animal models.[10] The research protocol adhered to the ethical guidelines approved by the University of Buea Institutional Animal Care and Use Committee (UB-IACUC), with permit number UB-IACUC # 04/2023, ensuring compliance with regulatory standards. Both male and female gerbils, aged 9-11 months, were randomly assigned to two groups: a control group (sham-operated) and an experimental group (implanted with *Onchocerca ochengi* worm masses). The worm masses used for implantation were obtained from infected cow skin purchased at a slaughterhouse in Ngaoundere, Cameroon. The specimens were transported under refrigeration to the University of Buea, where all experimental procedures were conducted.

### Isolation of *Onchocerca ochengi* Worm Masses

*O. ochengi* adult worm masses were isolated using the method described by Cho-Ngwa et al. (2010),[11] following established protocols for extracting parasites from infected host tissue. Briefly, fresh cattle skins with palpable nodules were thoroughly washed with tap water, rinsed with distilled water, and then towel-dried. After sterilization with 70% ethanol, the skins were left to dry in a laminar flow hood. The pale orange-yellow worm masses of *O. ochengi* were subsequently isolated. Worm masses were carefully excised using a razor blade to avoid damaging the adult worms. The masses were then immersed in incomplete culture medium (ICM) consisting of RPMI-1640, supplemented with 25 mM HEPES, 2 g/L sodium bicarbonate, 2 mM L-glutamine, 200 units/ml penicillin, 200 µg/ml streptomycin, and 0.25 µg/ml amphotericin B, adjusted to a pH of 7.4, in 24-well tissue culture plates. In some wells, worm embryos were observed while in other wells, motile microfilariae emerged from the adult worms and were also visible under the microscope. Thus worm masses implanted had adult females, males and microfilariae. Worm masses that were damaged or from decayed worms were excluded. Worms were confirmed for viability by MTT assay as before implantation in gerbils.

### Implantation of Worm Masses into Gerbils

To anesthetize the gerbils, a mixture of ketamine and diazepam (100 mg/kg and 5 mg/kg, respectively) was administered intraperitoneally using a 1 mL syringe equipped with a 25G needle. Once the animals were fully sedated, the abdominal area was shaved, and a small incision was made using a surgical blade. Test group animals were implanted with 10 or 15 *O. ochengi* worm masses, while control group animals underwent the same procedure without implantation. Nylon monofilament sutures were used to close the peritoneal membrane and abdominal wall separately to prevent worm displacement. The surgical area was sterilized with 70% ethanol, and the animals were then allowed to recover in cages with sterilized bedding. Post-operative care and monitoring were conducted daily to ensure recovery and health status.

### Behavioral Assessment

#### Acclimatization to Apparatus

On day 14 post-implantation, animals were introduced to the behavioral apparatuses—open field, elevated plus maze, and hanging wire—for familiarization. On day 18, animals were familiarized with the objects used for object recognition test. Each animal spent at least five minutes exploring each setup prior to the actual behavioral tests.

#### Elevated Plus Maze Test

The elevated plus maze test was performed to assess exploratory and emotional behaviour.[12] The apparatus, elevated 35 cm above the ground, consisted of two open and two closed arms. Gerbils were placed in the central platform, and their activity was observed for 5 minutes. Key behaviors recorded included time spent in the open and closed arms, frequency of entries into each arm, grooming, and head dips over the edges of the open arms.

#### Open Field Test

To assess anxiety and locomotion, gerbils were subjected to an open field test in a locally fabricated setup (60 cm x 60 cm area with 16 grid divisions, enclosed by 40 cm high walls). Each animal was placed at the center of the apparatus and observed for 5 minutes. The following behaviors were recorded: time spent in the center, number of grid crossings, defecation, urination, rearing, grooming, and stretch-attend posture.[13]

#### Object Recognition Test

The test was used to evaluate memory. On day 18, gerbils were introduced to the test apparatus with two identical objects and allowed to explore for 5 minutes. On day 19, one of the objects was replaced with a novel one, and exploration times for the familiar (tA) and novel (tB) objects were recorded. The discrimination between the previous and new object is a measure of memory performance.[14]

#### Hanging Wire Test

To evaluate neuromuscular function, the hanging wire test was conducted following the Treat-NMD Neuromuscular Network protocols. A wire was tensioned between two fixed stands, and animals were allowed to grasp the wire with their forelimbs. The time they could hold on before falling was recorded over three trials. A soft bedding area 35 cm below the wire prevented injury from falls.[15]

#### Sacrifice of Animals and Analysis

On day 21, the animals were re-sedated using the ketamine/diazepam combination, followed by sacrifice via cervical dislocation. Organs, including the brain, heart, liver, kidneys, lungs, and spleen, were extracted, weighed, and preserved for further analysis.

#### Recovery of Adult Worms

The abdominal cavity of each animal was dissected to recover implanted *O. ochengi* masses. Nodules were collected and incubated in 0.25 mg/mL liberase solution at 37 °C for one hour. Following incubation, worm viability was assessed using the MTT assay, which measures metabolic activity based on the conversion of MTT to formazan (purple crystals) by viable cells. Worms were scored for viability following the guide: 0% (no blue coloration seen), 25% inhibition of formazan, 50% inhibition of formazan, 75% inhibition of formazan, 90% inhibition of formazan to 100% (entire worm mass appeared blue) as previously reported[16].

#### Statistical Analysis

Data were processed using Microsoft Excel (2013) and analyzed using GraphPad Prism (version 8.4.3). Survival analysis was performed using the Kaplan-Meier method, with differences assessed via the Log-rank test. Comparisons between control and experimental groups were made using the Mann-Whitney U test. Statistical significance was set at P < 0.05. Data are presented as mean ± SEM.

## Results

### Physiological Effects of *O. ochengi* Worm Masses in Gerbils

The implantation of 15 *O. ochengi* worm masses per gerbil resulted in rapid mortality, with 90% of the test animals deceased by day 4 and 100% mortality by day 10 (Figure 1A). When 10 worm masses were implanted, 7 out of 15 animals died by day 10, with 5 gerbils succumbing by day 2 post-implantation (Figure 1B). None of the control animals died. A significant decrease in body weight was observed in the test animals by day 21 compared to the control group (Table 1). Among the organs weighed, only the spleen showed a significant difference, with the spleen of the test animals being three times heavier than that of the controls (Table 1). These findings indicate severe effects of *O. ochengi* worm masses on gerbil survival, body weight, and splenomegaly.

**Figure 1.**
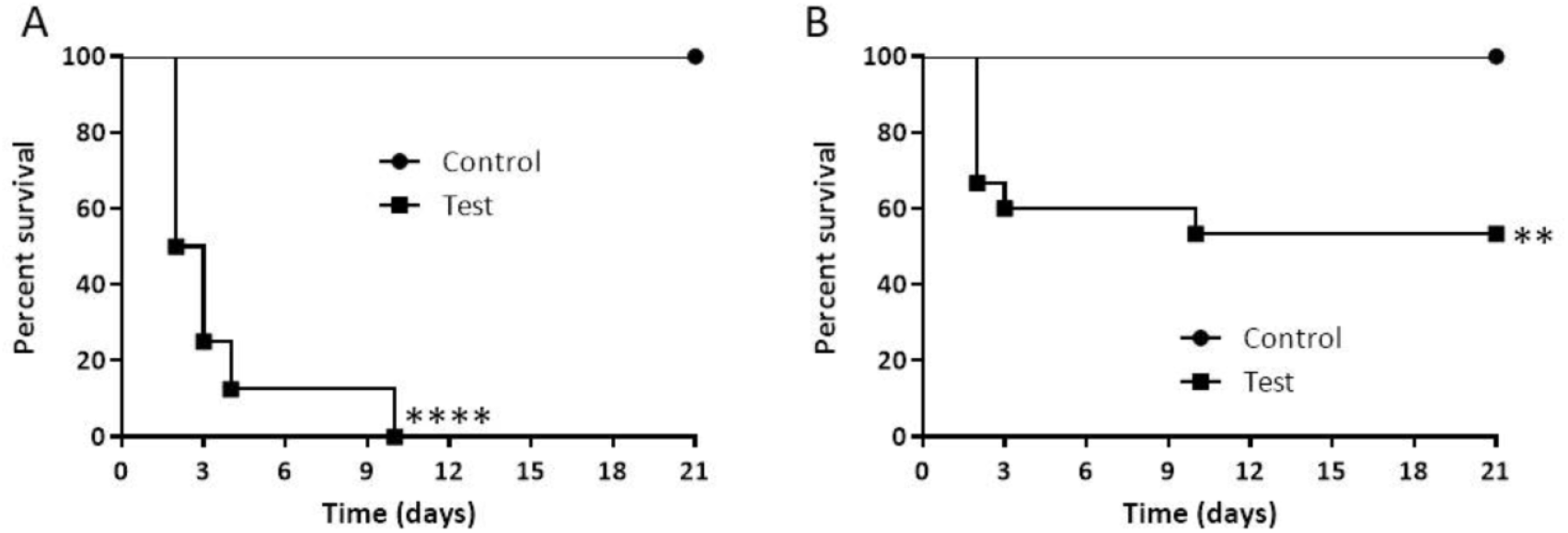
*O. ochengi* worm masses significantly decrease the survival of gerbils. Gerbil were surgically implanted with *O. ochengi* worm masses in the peritoneum (test group) or sham-operated (control group) and monitored for survival for 21 days: (A) Survival following implantation of 15 worm masses; n=8/group, (B) Survival following implantation of 10 worm masses; n=15/group. ****P<0.0001, **P=0.0028

**Table 1.** Gerbil percentage body weight and organ weight index on the day of sacrifice. Percentage body weight was obtained using animal weight on day 0 as baseline (100%) and organ index is the quotient of organ weight and total body weight on day 21. Control (n=10) and test (n=4). Data shows mean ± SEM

| Parameter | Organ | Control group | Test group | P value |
| --- | --- | --- | --- | --- |
| % body weight | | 99.7217±1.0151 | 92.4248±1.8787 | 0.0140* |
| Organ weight index | Liver | 0.0313±0.0016 | 0.0347±0.0029 | 0.1879 |
| | Spleen | 0.0008±0.0001 | 0.0024±0.0005 | 0.0080** |
| | Kidney | 0.0071±0.0002 | 0.0073±0.0005 | 0.9201 |
| | Eyes | 0.0059±0.0002 | 0.0061±0.0003 | 0.7333 |
| | Brain | 0.0158±0.0004 | 0.0167±0.0003 | 0.1419 |
| | Heart | 0.0043±0.0002 | 0.0042±0.0002 | >0.9999 |
| | Lungs | 0.0049±0.0003 | 0.0051±0.0001 | 0.3037 |

### Recovery of *O. ochengi* Worm Masses and Viability

Of the 8 surviving animals, 2 had no viable worms by day 21. Liberase digestion of worm mass nodules revealed numerous fragments of dead worms. Between 1 and 3 intact worms were recovered from the remaining 6 animals, with an average of 1.4 worms retrieved per 10 worm masses implanted (Table 2). The worms that were recovered displayed high viability, as shown by the MTT assay, with intense dark blue coloration (Figure 2B) and an average viability score of 93.3% (Table 2). Thus, while most of the worms died, those that survived by day 21 remained metabolically active.

**Table 2.** Number of *O. ochengi* worm masses recovered from test gerbils and MTT score. Gerbils were implanted with 10 *O. ochengi* worm masses on day 0 and analysed on day 21 and worms recovered assessed for viability by MTT assay.

| S/N | Number of worm masses recovered | MTT Scoring/% |
| --- | --- | --- |
| 1 | 2 | 90 |
| 2 | 1 | 85 |
| 3 | 3 | 100 |
| 4 | 0 | - |
| 5 | 2 | 95 |
| 6 | 2 | 100 |
| 7 | 1 | 90 |
| 8 | 0 | - |
| Average | 1.4 | 93.3 |

**Figure 2.**
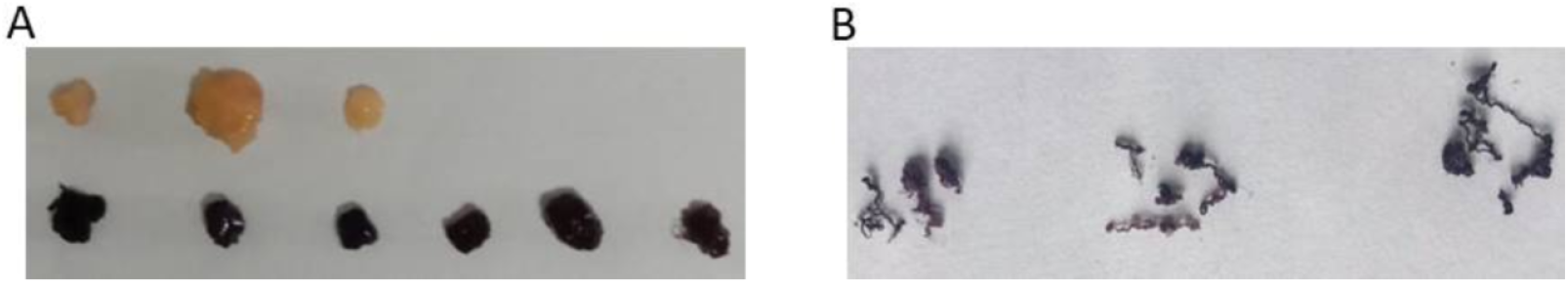
Adult worms after MTT assay. (A) Worms were assessed for viability before implantation; worms below were incubated with MTT while those above were not (negative control). (B) Picture of worms recovered from gerbils on day 21 after liberase treatment and MTT assay.

### Behavioral Assessment Results

The elevated plus maze test, used to evaluate emotional behavior and exploratory activity, showed no statistically significant differences between the test and control groups in any measured parameters (Table 3). Similarly, the open field test, which assesses anxiety and locomotion, did not reveal any significant differences between the groups across all 8 parameters (Table 3).

**Table 3.** No significant difference in elevated plus maze and open field tests. On day 15, control (n=15) and test (n=8) gerbils were assessed in an open field for 5 minutes for various parameters. Data shows mean±SEM

| Test | Parameter | Control group | Test group | P value |
| --- | --- | --- | --- | --- |
| **Elevated plus maze** | Duration at the center | 46.8±8.8 | 34.9±9.8 | 0.3320 |
|  | Number of entries in open arms | 14.9±3.0 | 13.4±3.3 | 0.6807 |
|  | Duration in open arm (s) | 94.5±13.5 | 69.8±16.2 | 0.3009 |
|  | Number of entries in closed arms | 17.4±1.6 | 16.9±1.9 | 0.7158 |
|  | Duration in closed arm (s) | 92.9±10.8 | 110.3±10.2 | 0.3325 |
|  | Number of head dipping | 32.5±4.6 | 18.4±3.2 | 0.0623 |
|  | Number of grooming | 0.1±0.1 | 0.0±0.0 | >0.9999 |
|  | Duration of grooming (s) | 0.1±0.1 | 0.0±0.0 | >0.9999 |
| **Open field** | Duration at the center (s) | 8.3±1.5 | 7.3±2.1 | 0.3456 |
|  | Number of lines crossed | 175.1±15.1 | 188.3±19.0 | 0.7396 |
|  | Number of stretch attempts | 0.0±0.0 | 0.0±0.0 | >0.9999 |
|  | Number of grooming | 0.8±0.3 | 3.0±0.9 | 0.1472 |
|  | Number of rearing | 62.5±6.0 | 62.6±6.4 | 0.8870 |
|  | Duration of rearing (s) | 66.8±4.6 | 58.0±8.1 | 0.2580 |
|  | Number of defecation | 3.1±0.9 | 2.9±0.3 | 0.5132 |
|  | Number of urination | 0.4±0.2 | 0.0±0.0 | 0.2507 |

In the object recognition test, designed to assess recognition memory, there was no significant difference in interaction time with old and new objects between the test and control groups (Table 4). The hanging wire test, used to measure muscle strength, showed no statistically significant difference, although there was a trend toward reduced muscle strength in the test group compared to the controls. Overall, none of the four behavioral assessment tests demonstrated statistically significant differences between the test and control groups.

**Table 4.**
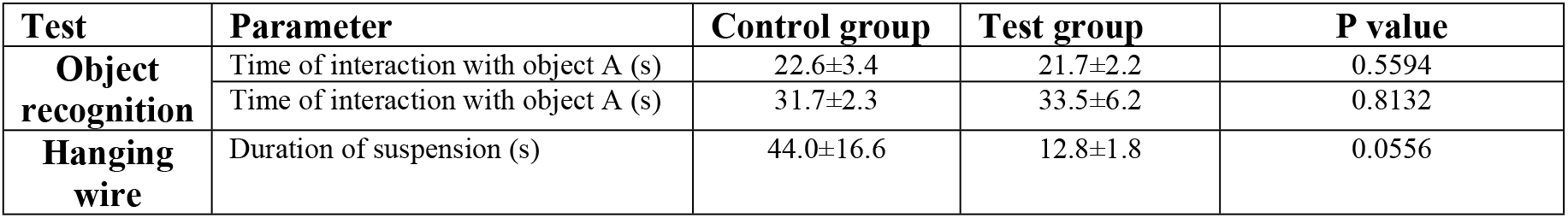
No significant difference in object recognition and hanging wire tests. Object recognition; control (n=10) and test (n=4) and hanging wire (n=5) and test (n=4); control Data represents mean ± SEM

## Discussion

The present study aimed to investigate the potential epileptogenic role of *O. ochengi* infection using a gerbil model. Despite previous epidemiological links between *Onchocerca* infections, particularly *O. volvulus*, and epilepsy in humans,[4, 6] no direct experimental evidence has supported a causal relationship between the parasitic infection and epilepsy. The study’s findings offer valuable insights into the physiological effects of *O. ochengi* infection, particularly in terms of mortality, weight loss, and organ pathology, but fail to provide conclusive evidence that the infection directly induces epilepsy-like symptoms in gerbils.

The marked increase in mortality rates observed in the gerbils implanted with *O. ochengi* worm masses is a clear indication of the severe systemic effects of the parasite. The rapid death of all animals implanted with 15 worm masses and more than half of those implanted with 10 masses underscores the pathogenic potential of *O. ochengi*. This aligns with the general understanding that parasitic infections can exert significant strain on host physiology, particularly in cases of high parasite burden.[17] The profound weight loss and splenomegaly observed in the test animals further highlight the systemic impact of the infection. Splenomegaly is commonly associated with immune responses to parasitic infections, indicating a heightened inflammatory state.[18, 19] These findings are consistent with prior studies that have documented similar physiological effects in hosts infected with parasitic nematodes.[20]

However, despite the clear physical impact of *O. ochengi* infection, no significant behavioral changes were observed in the infected gerbils that would suggest the development of epilepsy. The absence of differences between test and control groups in anxiety-related, locomotor, cognitive, and neuromuscular tests indicates that *O. ochengi* does not induce epilepsy-like symptoms in this model. This contrasts with the epidemiological evidence suggesting a potential link between onchocerciasis and epilepsy in humans.[2] One possible explanation is that the mechanisms driving epilepsy in humans with onchocerciasis may involve factors beyond the mere presence of the parasites, such as secondary inflammatory responses, immune-mediated damage, or neurotoxicity induced by parasite-related antigens.[21]

Furthermore, the behavioral tests employed in this study, such as the elevated plus maze and open-field tests, were primarily designed to detect anxiety and exploratory behavior, which may not fully capture the subtle neurological changes associated with epileptogenesis. While cognitive impairment and anxiety are common comorbidities of epilepsy,[22] the specific pathophysiological changes underlying seizure activity may not have been adequately assessed by the tests used in this study. It is also possible that the parasitic infection failed to reach the central nervous system (CNS), which would be necessary to trigger epilepsy. Given that *O. ochengi* primarily affects skin and subcutaneous tissues in its natural hosts,[23] it is conceivable that the parasite did not induce the same neuroinflammatory responses in gerbils as might occur in human infections with *O. volvulus*.

The lack of evidence linking *O. ochengi* infection to epilepsy in gerbils has important implications for understanding the pathophysiology of OAE. Previous studies have proposed several mechanisms by which *O. volvulus* might contribute to epilepsy, including neuroinflammation, blood-brain barrier disruption, and autoimmunity.[6] However, the current findings suggest that infection alone may not be sufficient to trigger epileptogenesis, at least in this animal model. This point to the possibility that OAE may involve a multifactorial process in humans, where the parasitic infection interacts with other risk factors, such as genetic predisposition, malnutrition, or co-infections, to produce the observed increase in epilepsy prevalence in endemic regions.[24, 25]

Given that the gerbil model did not exhibit any significant neurological symptoms, it remains unclear whether *O. ochengi* or other *Onchocerca* species can induce epilepsy under certain conditions or in other species. Moreover, *O. ochengi* has not been shown to induce epilepsy in its natural bovine host, but it remains the closest in phylogeny to *O. volvulus*[26] that can be used for an animal model, since *O. volvulus* do not survive in rodents.[27] Future studies should consider investigating the effects of *O. ochengi* or *O. volvulus* in other animal models more prone to CNS involvement, or use alternative methods, such as direct inoculation of parasite antigens into the brain, to assess their potential epileptogenic effects.

This study presents several limitations that may have affected the outcomes. First, the choice of the gerbil model, while useful for its susceptibility to seizure activity[7], may not accurately mimic the pathophysiology of onchocerciasis in humans. Gerbils are not natural hosts for *O. ochengi*, and thus the parasite may not exhibit the same tissue tropism or immune evasion strategies as in its natural bovine or human hosts. Additionally, the implantation of worm masses into the abdominal cavity, rather than allowing for a natural infection route, may not accurately reflect the disease progression seen in human onchocerciasis, where parasites reside primarily in the skin and subcutaneous tissues. Another important limitation is the relatively short duration of the study. Epileptogenesis is often a slow process that can take weeks or months to develop,[9] and it is possible that the 21-day observation period was insufficient to detect long-term neurological changes associated with epilepsy. Longer studies would be needed to fully assess whether chronic infection with *O. ochengi* could eventually lead to seizure activity in gerbils. One significant limitation affecting the interpretation of this study’s findings is the high mortality rate, which reduced the test group’s sample size for behavioral studies to only four animals. This raises the possibility that the gerbils that died may have been more prone to epileptogenic activity, potentially influencing the experimental outcomes. Nevertheless, the data collected from the surviving gerbils was considered the most representative of live subjects under the study conditions. To address this limitation, future experiments should aim to include a larger sample size to improve statistical power and reliability. Additionally, given that the onset of seizures in OAE typically occurs during childhood or adolescence (ages 3 to 18 years), future studies should prioritize using younger animals to better model the developmental timeline of the condition.

The findings of this study, while not supportive of a direct causal link between *O. ochengi* infection and epilepsy, have important clinical implications. The absence of a clear epileptogenic effect suggests that interventions targeting *Onchocerca* infections alone may not suffice to reduce epilepsy prevalence in endemic regions. This underscores the need for comprehensive health strategies that address other contributing factors, such as co-infections and malnutrition, alongside parasitic control measures. However, the severe physiological effects observed in the infected animals highlight the importance of continued efforts to control onchocerciasis through mass drug administration programs, as untreated infections can still cause significant morbidity. In terms of future directions, more detailed investigations into the immune and inflammatory responses triggered by *Onchocerca* infections may provide insights into the potential mechanisms linking onchocerciasis and epilepsy. Studies should also consider exploring the role of microfilariae, which are thought to play a role in CNS invasion, as well as the effects of co-infections with other neurotropic parasites, such as *Toxoplasma* or *Plasmodium*, which may synergistically contribute to the development of epilepsy in endemic regions.[28, 29] Lastly, human-based studies, such as post-mortem analyses of brain tissue from epilepsy patients in onchocerciasis-endemic regions, may help clarify the extent to which *Onchocerca* species directly impact the CNS.

In conclusion, while this study does not confirm a direct link between *O. ochengi* infection and epilepsy, it opens the door for further research into the complex interactions between parasitic infections and neurological disorders. In addition, the high mortality rate and limited observation period restrict the interpretation of these findings. Understanding the precise mechanisms at play is crucial for developing effective interventions to address epilepsy in onchocerciasis-endemic regions. To the best of our knowledge, this study represents the first attempt to establish an animal model for OAE.

## Acknowledgements

We are grateful to all the members of the Drug and Molecular Diagnostic (DMD) Laboratory, Biotechnology Unit, University of Buea, for all their contributions towards the realisation of this work.

## References

1. Thurman DJ, Beghi E, Begley CE, Berg AT, Buchhalter JR, Ding D, et al. Standards for epidemiologic studies and surveillance of epilepsy. Epilepsia 2012 Sep;52 Suppl 7:2–26.

2. Colebunders R, Hendy A, van Oijen M. Nodding Syndrome in Onchocerciasis Endemic Areas. Trends Parasitol 2016 Aug;32(8):581–3.

3. Murdoch ME, Asuzu MC, Hagan M, Makunde WH, Ngoumou P, Ogbuagu KF, et al. Onchocerciasis: the clinical and epidemiological burden of skin disease in Africa. Ann Trop Med Parasitol 2002 Apr;96(3):283–96.

4. Chesnais CB, Nana-Djeunga HC, Njamnshi AK, Lenou-Nanga CG, Boullé C, Bissek AZ, et al. The temporal relationship between onchocerciasis and epilepsy: a population-based cohort study. Lancet Infect Dis 2018 Nov;18(11):1278–86.

5. Quek S, Hadermann A, Wu Y, De Coninck L, Hegde S, Boucher JR, et al. Diverse RNA viruses of parasitic nematodes can elicit antibody responses in vertebrate hosts. Nature Microbiology 2024;9(10):2488–505.

6. Hadermann A, Amaral LJ, Van Cutsem G, Siewe Fodjo JN, Colebunders R. Onchocerciasis-associated epilepsy: an update and future perspectives. Trends Parasitol 2023 Feb;39(2):126–38.

7. Loskota WJ, Lomax P, Rich ST. The gerbil as a model for the study of the epilepsies. Seizure patterns and ontogenesis. Epilepsia 1974 Mar;15(1):109–19.

8. Morales-Hojas R, Cheke RA, Post RJ. A preliminary analysis of the population genetics and molecular phylogenetics of Onchocerca volvulus (Nematoda: Filarioidea) using nuclear ribosomal second internal transcribed spacer sequences. Mem Inst Oswaldo Cruz 2007 Nov;102(7):879–82.

9. Pitkänen A, Lukasiuk K, Dudek FE, Staley KJ. Epileptogenesis. Cold Spring Harb Perspect Med 2015 Sep 18;5(10).

10. Ayiseh RB, Mbah GE, Manfo FPT, Kulu TK, Njotu FN, Monya E, et al. Survival of worm masses of Onchocerca ochengi in gerbils and hamsters: implications for the development of an in vivo macrofilaricide screening model. Parasitol Res 2023 Jul;122(7):1581–91.

11. Cho-Ngwa F, Abongwa M, Ngemenya MN, Nyongbela KD. Selective activity of extracts of Margaritaria discoidea and Homalium africanum on Onchocerca ochengi. BMC complementary and alternative medicine 2010;10:62.

12. Varty GB, Morgan CA, Cohen-Williams ME, Coffin VL, Carey GJ. The gerbil elevated plusmaze I: behavioral characterization and pharmacological validation. Neuropsychopharmacology 2002 Sep;27(3):357–70.

13. Belovicova K, Bogi E, Csatlosova K, Dubovicky M. Animal tests for anxiety-like and depression-like behavior in rats. Interdiscip Toxicol 2017 Sep;10(1):40–3.

14. Leger M, Quiedeville A, Bouet V, Haelewyn B, Boulouard M, Schumann-Bard P, et al. Object recognition test in mice. Nat Protoc 2013 Dec;8(12):2531–7.

15. van Putten M, de Winter C, van Roon-Mom W, van Ommen GJ, t Hoen PA, Aartsma-Rus A. A 3 months mild functional test regime does not affect disease parameters in young mdx mice. Neuromuscul Disord 2010 Apr;20(4):273–80.

16. Bulman CA, Bidlow CM, Lustigman S, Cho-Ngwa F, Williams D, Rascón AA, Jr., et al. Repurposing auranofin as a lead candidate for treatment of lymphatic filariasis and onchocerciasis. PLoS Negl Trop Dis 2015 Feb;9(2):e0003534.

17. Brattig NW. Pathogenesis and host responses in human onchocerciasis: impact of Onchocerca filariae and Wolbachia endobacteria. Microbes Infect 2004 Jan;6(1):113–28.

18. Vincent AL, Ash LR. Splenomegaly in Jirds (Meriones Unguiculatus) Infected with Brugia Malayi (Nematoda: Filarioidea) and Related Species. The American Journal of Tropical Medicine and Hygiene 1978 01 May. 1978;27(3):514–20.

19. Gharbi HA, Marbot P, Cheikh MB. Splenic Involvement in Parasitoses. Ultrasonography of the Spleen. Berlin, Heidelberg: Springer Berlin Heidelberg; 1988. p. 70–85.

20. Bethony J, Brooker S, Albonico M, Geiger SM, Loukas A, Diemert D, et al. Soil-transmitted helminth infections: ascariasis, trichuriasis, and hookworm. Lancet 2006 May 6;367(9521):1521–32.

21. Galán-Puchades MT. Onchocerciasis-associated epilepsy. The Lancet Infectious Diseases 2019 2024/10/18;19(1):21–2.

22. Fisher RS, Acevedo C, Arzimanoglou A, Bogacz A, Cross JH, Elger CE, et al. ILAE Official Report: A practical clinical definition of epilepsy. Epilepsia 2014;55(4):475–82.

23. Allen JE, Adjei O, Bain O, Hoerauf A, Hoffmann WH, Makepeace BL, et al. Of Mice, Cattle, and Humans: The Immunology and Treatment of River Blindness. PLOS Neglected Tropical Diseases 2008;2(4):e217.

24. Olum S, Scolding P, Hardy C, Obol J, Scolding NJ. Nodding syndrome: a concise review. Brain Commun 2020;2(1):fcaa037.

25. Benedek G, Abed El Latif M, Miller K, Rivkin M, Ramadhan Lasu AA, Riek LP, et al. Protection or susceptibility to devastating childhood epilepsy: Nodding Syndrome associates with immunogenetic fingerprints in the HLA binding groove. PLoS Negl Trop Dis 2020 Jul;14(7):e0008436.

26. Morales-Hojas R, Cheke RA, Post RJ. Molecular systematics of five Onchocerca species (Nematoda: Filarioidea) including the human parasite, O. volvulus, suggest sympatric speciation. J Helminthol 2006 Sep;80(3):281–90.

27. Allen JE, Adjei O, Bain O, Hoerauf A, Hoffmann WH, Makepeace BL, et al. Of Mice, Cattle, and Humans: The Immunology and Treatment of River Blindness. PLoS neglected tropical diseases 2008;2(4):e217.

28. Ngoungou EB, Bhalla D, Nzoghe A, Dardé ML, Preux PM. Toxoplasmosis and epilepsy--systematic review and meta analysis. PLoS Negl Trop Dis 2015 Feb;9(2):e0003525.

29. Trivedi S, Chakravarty A. Neurological Complications of Malaria. Curr Neurol Neurosci Rep 2022 Aug;22(8):499–513.

